# Global diversification of the common moonwort ferns (*Botrychium lunaria* group, Ophioglossaceae) was mainly driven by Pleistocene climatic shifts

**DOI:** 10.1101/2022.09.28.509846

**Authors:** Vinciane Mossion, Erik Koenen, Jason Grant, Daniel Croll, Donald R. Farrar, Michael Kessler

**Affiliations:** Laboratory of Evolutionary Genetics, University of Neuchâtel, Neuchâtel, Switzerland; Department of Systematic and Evolutionary Botany, University of Zürich, Zurich, Switzerland; Department of Ecology, Evolution and Organismal Biology, Iowa State University of Science and Technology, United-States of America

**Keywords:** Ophioglossaceae, ferns, speciation, phylogenetics, climate, ploidy

## Abstract

The cosmopolitan *Botrychium lunaria* group belong to the most species rich genus of the family Ophioglossaceae and was considered to consist of two species until molecular studies in North America and northern Europe led to the recognition of multiple new taxa. Recently, additional genetic lineages were found scattered in Europe, emphasizing our poor understanding of the global diversity of the *B. lunaria* group, while the processes involved in the diversification of the group remain unexplored. We conducted the first global phylogenetic study of the group including 513 ingroup accessions sequenced for four non-coding plastid loci. We recovered ten well-supported clades, although relationships between clades were inconsistent between Bayesian and Maximum Likelihood analyses. We treated each clade at the species level, except for one clade including two ploidy levels, resulting in the recognition of 11 species, 5 of which are unnamed. In contrast to previous studies, we found species diversity to be equally distributed across the northern hemisphere, with 7-8 species per continent. We estimated the stem age of the *B. lunaria* group at 2.4-5.1 million years, with most species 1.8-2.6 million years old, and subspecies 0.5-1.0 million years old. Diversification thus coincided with Pleistocene climatic fluctuations that strongly affected the areas inhabited by the group, suggesting that diversification was driven by climatically induced cycles of extinction, dispersal, and migration. Furthermore, ecological differentiation between species suggests these complex population dynamics were associated with adaptations to specific environmental conditions. We found limited evidence that speciation is driven by polyploidization and hybridization. We show that the *B. lunaria* group has greater species level diversity than previously assumed and suspect that further cryptic species may await discovery, especially in the *B. neolunaria* clade.

## Introduction

Ferns are an ancient group of vascular plants, with a fossil records dating back to the middle Devonian (387.7 – 382.7 million years) (Berry and Hilton, 2006; Taylor et al., 2009) and divergence from the seed plants estimated to have taken place about 360-430 million years (Des Marais et al., 2003; Lehtonen et al., 2017; Magallón et al., 2013; Pryer et al., 2004; Qi et al., 2018; Rothfels et al., 2015; Testo and Sundue, 2016; Wikström and Kenrick, 2001; Zhong et al., 2014). Devonian fern lineages such as Cladoxylopsida or Rhacophytales (Taylor et al., 2009) have long gone extinct, while most of the about 11,000 extant fern species arose from a Cenozoic radiation (Schneider et al., 2004; Schuettpelz and Pryer, 2009; Testo and Sundue, 2016). The recent nature of most extant fern species and their overall slow (Smith, 1972), but heterogeneous (Rothfels et al., 2012; Testo and Sundue, 2016) rates of diversification, imply that many groups comprise species complexes characterized by morphologically poorly-defined species (Paris et al., 1989; Vasques et al., 2019; Williams et al., 2016), often in combination with polyploid networks in which diploid species give rise to allopolyploid hybrids which then in turn diversify (Chang et al., 2013; Hanušová et al., 2019; Lovis, 1958; Rothfels et al., 2014; Williams et al., 2016). In such a context, interpreting species diversity and understanding the speciation processes involved in diversification is challenging and intriguing.

The Ophioglossaceae is an example of an early diverging family of ferns estimated at 262-322 million years old (Kumar et al., 2017; Lehtonen et al., 2017; Testo and Sundue, 2016), composed of recently diverged species (Dauphin et al., 2018; Lehtonen et al., 2017; Rothfels et al., 2015) many of which are genetically distinct but morphologically hardly distinguishable (Dauphin et al., 2017; Hauk, 1995; Zhang et al., 2020). Ophioglossaceae are characterized by subterranean gametophytes that are colonized by mycorrhizal fungi (Winther and Friedman, 2007), and an unusual sporophyte leaf morphology: the single leaf produced each year is composed of a sterile photosynthetic blade, the trophophore, and of a fertile portion, the sporophore (Figure 1A). This simple morphology provides only a limited number of distinctive features, making morphological differentiation of species challenging (Farrar, 2011; Stensvold, 2008; Williams, 2015; Williams et al., 2016). Accordingly, the number of species recognized in the family has historically been quite low. Clausen (1938) only recognized six species in *Botrychium sensu stricto* (*subgenus Eubotrychium*). Later Wagner (Wagner and Grant, 2002; Wagner and Wagner, 1981, 1994, 1990a, 1990b, 1986, 1983a, 1982) and other taxonomists (Farrar and Johnson-Groh, 1991, 1991; Gilman et al., 2015; Mickel and Smith, 2004; Popovich et al., 2020; Stensvold et al., 2002; Stensvold and Farrar, 2017), utilizing cytology, allozyme data, and detailed morphological analyses gradually increased the number of taxa to currently 36 (Dauphin et al., 2017; PPG I, 2016), of which 30 have been described in the last four decades. The newly defined species exhibit subtle morphological differences often along with geographical and ecological distinctness. More recently, phylogenetic analysis based on non-coding plastid markers (rbcL, trnL L-F, rpl16 intron, psbA-trnH^GUG^, matK intron) (Dauphin et al., 2017, 2014; Hauk, 1995; Hauk et al., 2012, 2003) uncovered further lineage diversity within *Botrychium*, especially among the European populations treated as *Botrychium lunaria* (L.) Sw. (Dauphin et al., 2017, 2014; Maccagni et al., 2017). These findings suggested that deeper investigations on *B. lunaria* populations might shed further light on the diversity within the group.

**Figure 1:**
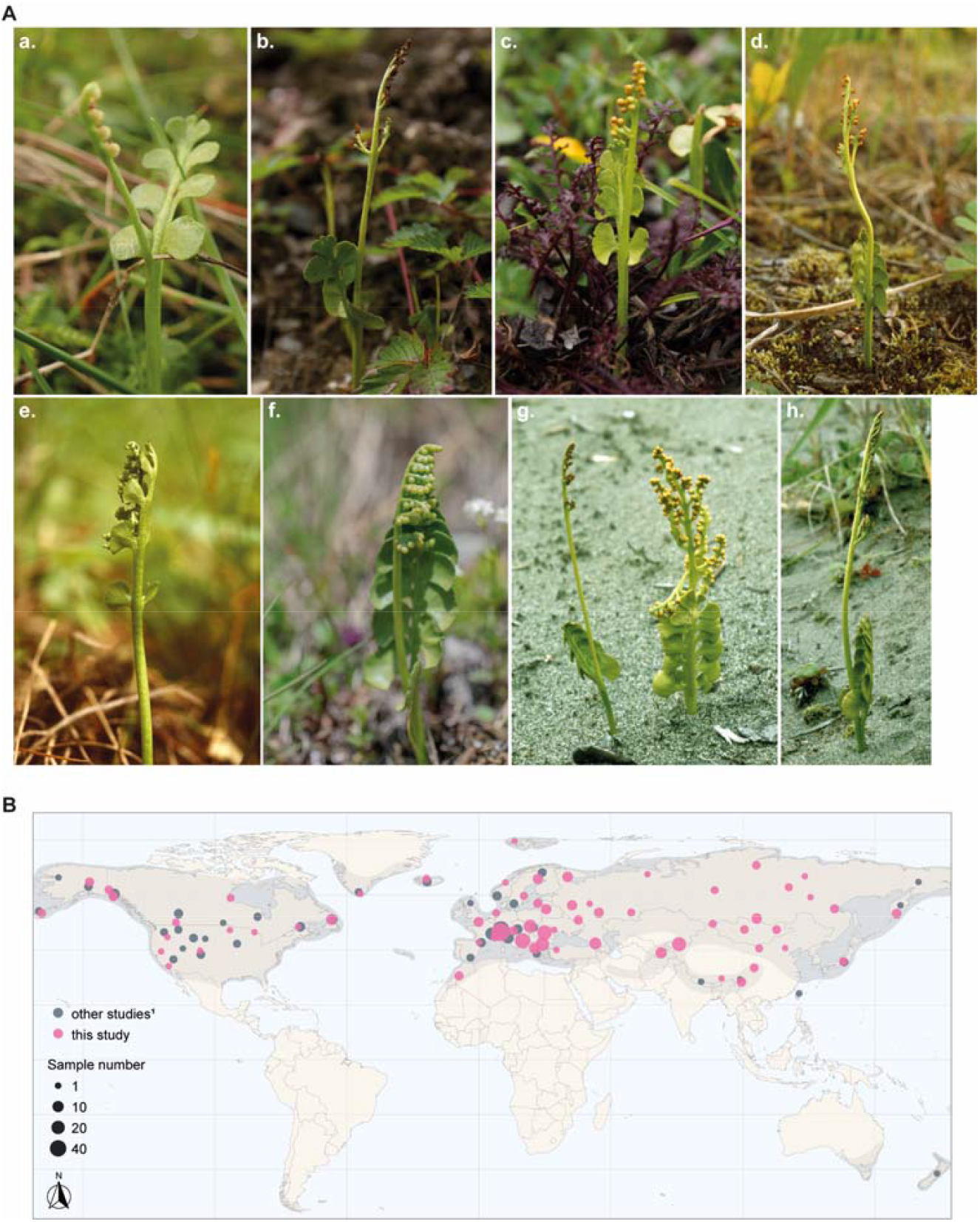
Morphological diversity of the B. lunaria group and geographical distribution of the analyzed individuals. (A) **a**. VM-lun26-5-RO (photo by Q. Dubois, Romania). **b**. VM-lun55-1-CN (photo by V. Mossion, China). **c**. VM-lun54-CN (photo by V. Mossion, China). **d**. *B. neolunaria* (photo by D. Farrar, USA). **e**. *B. crenulatum* (photo by D. Farrar, USA). **f**. *B*. lunaria, VM-lun89-CH (photo by V. Mossion, Switzerland). **g**. *B. neolunaria s*.*str*. on the left, *B. tunux* on the right (photo by D. Farrar, USA). **h**. *B. yaaxudakeit* (photo by D. Farrar, USA). (B) Sampling map, differentiating between sequences from previous studies (gray) and this study (pink). The distribution range of the *B. lunaria* group (Stensvold, 2018) is showed by the grey layer. Individuals less than 500 km apart were clustered and are indicated by a unique circle. The circle sizes are proportional to individual number. ^1^Dauphin *et al*. (2014, 2017), Maccagni *et al*. (2017).

The *Botrychium lunaria* group is one of three monophyletic clades within *Botrychium*, called the Lunaria clade by (Dauphin et al., 2014). It includes what was long considered to be a single, morphologically variable species that occurs across the boreal and alpine regions of North America, Europe, and Asia with outposts in New Zealand and Australia (Clausen, 1938; Kato, 1987; Milde, 1869). As early as 1903, Underwood suggested that at least two distinct taxa occurred in North America, prompting his description of *B. onondagense*. However, subsequent studies treated *B. onondagense* as a variety or form of a *B. lunaria* (Butters and Abbe, 1953; Clarke and House, 1923; Clausen, 1938; Clute, 1905). Later, based on extensive morphological surveys, Wagner (1981) described *B. crenulatum*, a North American species, which has been widely accepted (Dauphin et al., 2017; Farrar et al., 2017; Stensvold and Farrar, 2017). When allozyme markers were used more extensively and the investigation area was extended to northern Europe, five additional taxa were recognized (*B. tunux, B. nordicum, B. neolunaria, B. yaaxudakeit*, and *B. lunaria* var. *melzeri*) (Stensvold et al., 2002; Stensvold and Farrar, 2017). The recognition of *B. neolunaria* as the common *B*. “*lunaria*” in North America re-evaluated Underwood’s *B*. “*onondagense*” as the North American representative of European *B. “lunaria*”. Until a few years ago, most taxa in the *B. lunaria* group were from in North America. More recently, however, it has been found that the genetic diversity among *B. “lunaria”* populations in western and central Europe may exceed the diversity found in North America (Dauphin et al., 2014, 2014; Maccagni et al., 2017). Some European lineages could be assigned to species previously described from North America, such as *B. tunux* (Stensvold et al., 2002). Yet, there remain several poorly understood lineages within *Botrychium* possibly representing additional species (Dauphin et al., 2017). These hitherto undescribed lineages are widespread throughout Europe, highlighting the uncertainty of taxonomic assignments of European *B. “lunaria”* and the limited understanding of the global diversity of the *B. lunaria* group.

Significant advances have been made to understand speciation processes within *Botrychium*. Polyploidization appears to be a major factor in some clades of *Botrychium*, and complex polyploid networks have been uncovered (Dauphin et al., 2016; Wagner, 1993; Wagner and Lord, 1956; Williams and Waller, 2012), but the relevance of polyploidization in the *B. lunaria* group remains unclear. Among the 33 accepted *Botrychium* species, nearly half are allopolyploids. Of the new taxa in the *B. lunaria* group, *B. yaaxudakeit* is noteworthy as the first and only allopolyploid species discovered in the group. Using allozyme analysis, Stensvold *et al*. (2002) found that *B. yaaxudakeit* carries a combination of alleles matching *B. “lunaria”* and *B. neolunaria* and shows morphological intermediacy. These criteria support the hypothesis that *B. yaaxudakeit* was derived by hybridization between the two diploid species followed by chromosome doubling. Scattered sampling in North America and central Europe (Dauphin et al., 2016; Veselý et al., 2012; Williams and Waller, 2012) found no further evidence of polyploids in the *B. lunaria* group, suggesting that the speciation processes involved in the diversification of this group may differ from other *Botrychium* clades. However, diploid hybrids between *B. neolunaria* and *B. “lunaria”* have been detected by allozyme markers in North America, in western and northern Europe, and on the east coastline of Asia and Oceania (Stensvold et al., 2002; Stensvold and Farrar, 2017). These hybrids, commonly called introgressed hybrids, originate from the germination of spores produced by F1 hybrids. The term reflects the unequal contributions by the two diploid parents during the meiosis. Introgressed hybrids are diploid and fertile plants which are supposedly as isolated from their parents as allotetraploids are. The success of the hybrids suggests that introgression may favor rapid speciation by promoting adaptive divergence (Abbott et al., 2013). Phylogenetic analysis of plastid data showed that whereas the maternal donor of these hybrids was *B. neolunaria*, hybrids formed a distinct subclade (Dauphin et al., 2017), supporting this idea. Hence, both polyploidization and hybridization events have contributed to the diversification within the *B. lunaria* group, but their specific contributions remain poorly understood.

Diversification within the *B. lunaria* group may have been driven by successive isolation, migration, and dispersal events imposed by the Quaternary glacial-interglacial cycles (Stensvold, 2008). The *B. lunaria* group mainly occurs in circumboreal-temperate regions strongly affected by Quaternary glacial cycles (Ehlers et al., 2011; Stensvold, 2008). As in many other boreal and alpine plant groups, this may have led to complex population dynamics (Schönswetter et al., 2005), possibly associated with adaptations to different climatic or soil conditions (Allen et al., 2012; Alvarez et al., 2009; Lafontaine et al., 2018). For some species of the genus *Botrychium*, local ecological, climatic, and edaphic specialization has been reported. For example, populations of *B. campestre* from northeastern Iowa (USA) have slope preferences within habitats (Nekola and Schlicht, 1996), whereas populations of *B. matricariifolium* from Vosges (France) occur on sandy, acid soils, in areas with high annual mean precipitation and cool summers (Muller, 1986). The species *B. simplex* and *B. tenebrosum*, which form a closely related species pair that occurs in North America, Sweden, and Switzerland, also tend to have different preferences for wetter or dryer habitats, respectively (Ståhl et al., 2016; A. Maccagni pers. comm.). However, these species belong to other clades of *Botrychium*, and little is known on the climatic and edaphic niches of species from the *B. lunaria* group. Ecological niche segregation among two genetic lineages of *B*. “*lunaria*” has been suggested based on habitat preferences in the Alps (Maccagni et al., 2017). In North America, *B. tunux* and *B. yaaxudakeit* favor well drained soils whereas *B. neolunaria* grows on poorly to moderately drained substrates (Stensvold et al., 2002; Stensvold and Farrar, 2017). In the absence of further characterization of the ecological niches of the lineages within the *B. lunaria* group, it is currently impossible to address to which degree climatic and edaphic adaptation contributed to the diversification of the group.

In this study, we analyzed the worldwide diversity and diversification time of the *B. lunaria* group and associated the occurrence of individual lineages with climatic and edaphic factors. We performed a plastid-based phylogenetic reconstruction including a large set of newly collected samples from eastern Europe and Asia. We also investigated genome size variation to identify potential polyploidization events throughout the diversification of the *B. lunaria* group. Specifically, we asked the following questions: 1) How many species-level taxa can be recognized in the *B. lunaria* group? 2) What is the species diversity of the group in eastern Europe and Asia? 3) What is the role of polyploidization in the diversification of the group? 4) Can the timing of species divergences can be related to Pleistocene climatic fluctuations? 5) Is there ecological differentiation between the species that may be linked to the diversification of the group?

## Material and Methods

### 1. Sample collection

We collected 172 fresh sporophytes at 63 locations within Eurasian Mountain ranges, including 41 sites in central and eastern Europe (Alps, Carpathians, Dinaric Alps, Rhodopes, Central Balkans, and Caucasus) and 14 in central and eastern Asia (Pamir-Alay, Tien-San, Eastern Himalayas, Qinling, and Yann) (Figure 1B). At each location, we sampled 3-15 plants at least 20 cm apart, collected soil samples around the roots of one or two individuals, measured the soil pH using a Hellige pH-meter and recorded the geographical coordinates at the center of the sampling site using a Garmin eTrex 30X GPS. From each plant, a few pinnae were silica-dried on site and the remaining plant kept as herbarium voucher.

Vouchers were deposited in the herbaria of the University of Zurich (Z/ZT) and University of Neuchâtel (NEU), Switzerland, and in herbaria in the countries of origin (supplementary Table S1). In addition, we also incorporated 203 samples from China, Russia, Europe, North Africa, and North America provided by collaborators (see acknowledgments) and herbaria (PE, MW, AIX, ISC, MARK) (supplementary Table S1), which included type material of five recognized taxa (*B. tunux, B. neolunaria, B. crenulatum, B. yaaxudakeit, B. lunaria* var. *melzeri*). The sampling was designed to cover the whole distribution range of the *B. lunaria* group as described by Stensvold (2008) (Figure 1B).

### 2. DNA extraction, amplification, and sequencing

We extracted total genomic DNA from silica-dried and herbarium material using the DNeasy plant mini kit (Qiagen, Hilden, Germany). We modified the manufacturer’s protocol as follows: the mixture was incubated in buffer AP1 and RNase A for 20 minutes, a centrifugation step of 1 minutes at 8000 rpm was added before the centrifugation of the 17^th^ step, and DNA was eluted in ultra-pure water. Total genomic DNA was quantified and quality-checked by spectrophotometry (NanoDrop 2000). The four non-coding plastid loci targeted (trnH^GUG^-psbA intergenic spacer, trnL^UAA^-trnF^GAA^ intergenic spacer, matK intron, and rpl16 intron) were amplified by polymerase chain reaction (PCR) following protocols and thermocycling conditions published by Dauphin *et al*. (2014; 2017) and Small *et al*. (2005). Besides, we optimized thermocycling conditions for touchdown protocol and the use of a high-fidelity DNA polymerase to amplify DNA degraded and extracted from herbarium material (supplementary Table S2). Briefly, the PCR reactions were performed in a volume of 50 µl containing 10 ng of DNA template, 0.2 µM of each primer (Microsynth, Balgach, Switzerland), 0.2 mM of DNTPs mix (Promega, Madison, USA), 1 unit of GoTaq DNA polymerase (Promega), 1X GoTaq buffer (Promega), and an additional 1 mM of MgCl_2_ (Promega) for trnL^UAA^-trnF^GAA^. Herbarium material DNAs were amplified using Q5 high-fidelity DNA polymerase with the following PCR product concentrations: 0.3 µM of each primer (Microsynth), 0.2 mM of DNTPs mix (Promega), 0.5 unit of Q5 (New England Biolabs Inc., Ipswich, England), 1X Q5 buffer (New England Biolabs Inc.), and an additional 1 mM of Q5 enhancer (New England Biolabs Inc.) for trnL^UAA^-trnF^GAA^. Each PCR product was quality-checked by electrophoresis on agarose gel (1.5 %) stained with GelRed® (Biotium, Fremont, USA) and visualized under UV light on a transilluminator (UVI Tec Gel Documentation System). Template purifications and sequencing were outsourced to Macrogen Europe (Amsterdam, Netherlands). Succinctly, sequencing reactions were performed in the Master Cycler pro 384 (Eppendorf) using the ABI BigDye® Terminator v3.1 Cycle Sequencing Kit (Applied Biosystems), following the protocols supplied by the manufacturer. Single-pass sequencing was performed on each template using the same primers combination than PCR. The fluorescent-labeled fragments were purified from the unincorporated terminators with the BigDye XTerminator® Purification Kit (Applied Biosystems). The samples were injected to electrophoresis in an ABI 3730xl DNA Analyzer (Applied Biosystems). The four regions were paired end sequenced. The remaining silica material and DNA extractions were deposited at the Botanical Institute of Zurich, Switzerland.

### 3. Sequence cleaning and alignment

Sequences were trimmed and ambiguous bases annotated using the default settings of Geneious v.8.1.9 (https://www.geneious.com) *Trim ends* and *Find heterozygotes* plug-ins. Consensus sequences were obtained from paired end contigs, and uncertain position were manually cleaned according to the base quality of the chromatograms. In the case of ambiguous bases presenting equal chromatogram quality on both sequences, ambiguity was kept, and the base coded according to the IUB code (Cornish-Bowden, 1985). Consensus sequences were aligned per locus using MAFFT v. 7.017 (Katoh and Standley, 2013) under the G-INS-i strategy and default parameters. Alignments were visually inspected and manually curated using Geneious. For instance, gaps in low quality 3’ and 5’ end regions were annotated as missing data. A minority of individuals presented partial sequencing data (6 %, supplementary Table S1, Table 1). Additional published sequences of *Botrychium* (*e*.*g*., outgroups, specimens of the synonymized *B. onondagense* and supplementary type material) and of the Botrychioideae sub-family were incorporated to our dataset (Dauphin et al., 2017, 2014; Maccagni et al., 2017; Zhang et al., 2020) (supplementary Table S1, S3). A third of these supplementary accessions had information at the trnH^GUG^-psbA and trnL^UAA^-trnF^GAA.^ loci only. The uniparentally inheritance (Gastony and Yatskievych, 1992; Guillon and Raquin, 2000; Kuo et al., 2018; Vogel et al., 1998) and the absence of recombination (Ravi et al., 2008) in fern chloroplasts allowed to analyze the four targeted loci as a concatenated single locus. Alignments per locus were concatenated using the R v.4.0.3 (R Core Team, 2018) seqinr v.4.2-4 (Charif and Lobry, 2007) package and the *concat* function (Khang, 2016). The ingroup was represented by 521 individuals with type material of each named taxon within the *B. lunaria* group except for *B. lunaria* and *B. onondagense*. The outgroup contained nine diploid species of the two other *Botrychium* clades (Lanceolatum and Simplex-Campestre) (supplementary Table S1).

**Table 1:**
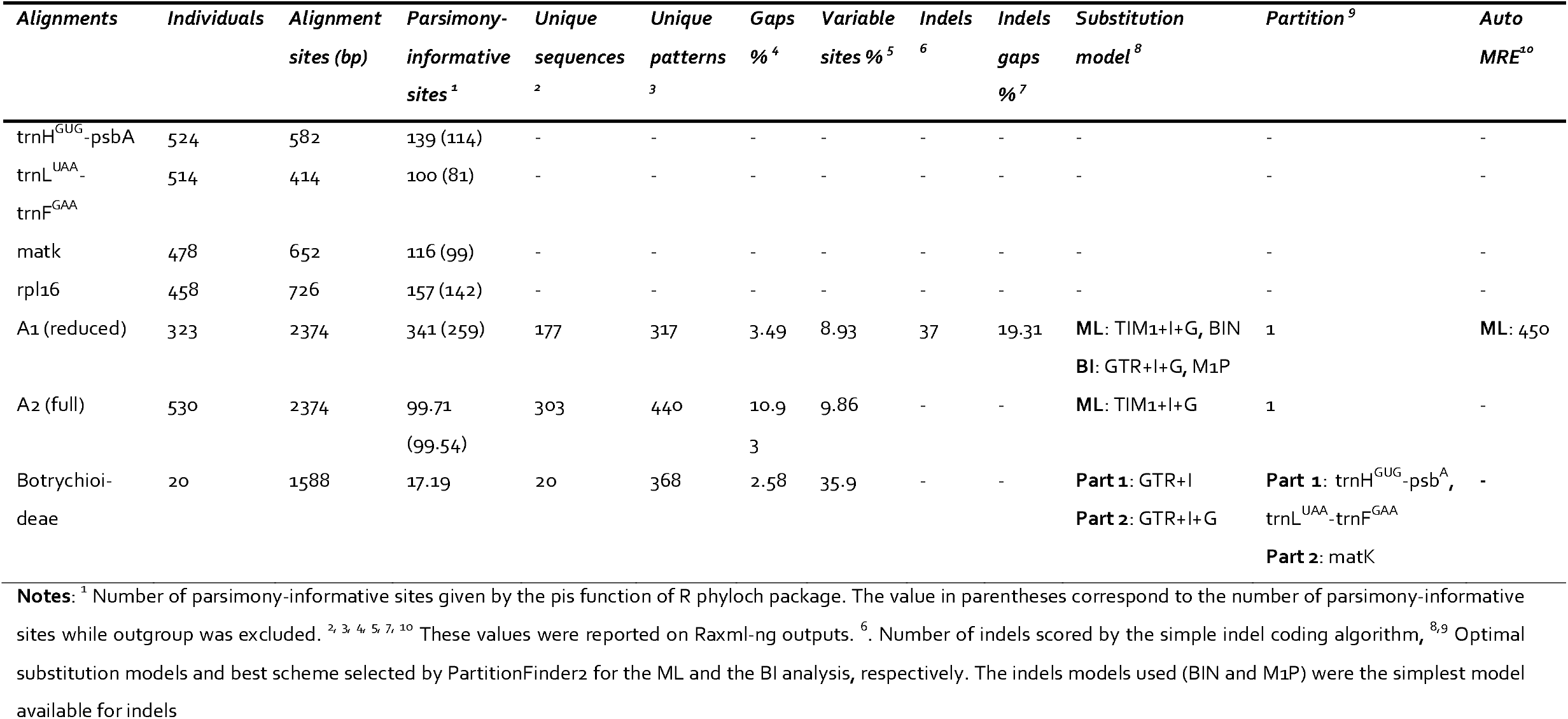
Characteristics of the multiple alignments including the outgroup.

### 4. Phylogenetic reconstruction

We assessed the phylogenetic relationships within the *B. lunaria* group based on two datasets (*i*.*e*., multiple alignment): one containing all the 530 accessions (full dataset - A2; supplementary File S1) and one containing a subset of 323 individuals that excluded 82 samples with partial sequencing data and 116 accessions carrying low phylogenetic signal (rogue individuals) (reduced dataset - A1; supplementary File S2). Rogues were identified using RogueNaRok (Aberer et al., 2013). The calculations were based on one hundred bootstrap trees estimated with the best tree inferenced from the multiple alignment exempt of partial data under a TIM1+I+G substitution model using RAxML-NG v. 1.0.1 (Kozlov et al., 2019) (Table 1). For each dataset, the best scheme and the optimal substitution model were assessed using PartitionFinder 2 (Frandsen et al., 2015) with *branch lengths* set as *unlinked*, and the *substitution models* set as *all* or *MrBayes*. Optimal substitution models were chosen according to the Akaike information criterion (AICc). Indels were scored for the A1 dataset using the *2 matrix* script (Salinas and Little, 2014) which implemented the simple indel coding algorithm described by (Simmons and Ochoterena, 2000). In total, 36 indels were scored (supplementary File S2). Unrooted species trees were built with maximum likelihood (ML) using RAxML-NG and Bayesian inferences (BI) using Bayesphylogenies 2 (Pagel et al., 2004).

The reduced dataset was analyzed both with and without indels. We ran ML inferences under a TIM1+I+G substitution model and BIN model for the indels with a fixed random seed of 2. The tree search was performed on 25 random and 25 parsimony-based starting trees and branch support was estimated over 1,000 bootstrap replicates. The minimum number of bootstrap replicates was also determined using the autoMRE bootstrap convergence test (Pattengale et al., 2010). Bootstrap values were depicted on the best tree using the *Transfer Bootstrap Expectation* metric (Lemoine et al., 2018). BI were conducted under GTR+I+G substitution model and M1P models for indels using the reversible jump method. We set two runs of 50 million MCMCMC generations, each including one cold and 2 heated chains, with a seed number of 946432 and tree sampled every 5,000 generations. Parameter files were inspected with tracer v. 1.7 (Rambaut et al., 2018)and a burn-in of 10 and 14% was applied to the analyses with and without indels, respectively. Majority-rule consensus (MRC) trees of 50% were built based on posterior probabilities (PP) using SumTrees v. 4.4.0 and DendroPy library v. 4.4.0 (Sukumaran and Holder, 2010). The full dataset was analyzed with a constraint backbone to minimize the effect of incomplete data and rogue individual on tree inference. We used the best ML tree of the reduced dataset as the constraint backbone. This phylogenetic inference was performed by ML only under a TIM1+I+G substitution model with a fixed random seed of 2. The tree search was performed on 25 random and 25 parsimony-based starting trees and branch support was estimated over 100 bootstrap replicates. Trees were depicted using R ggtree v. 2.4.0 (Yu et al., 2018, 2017), ape v. 5.4-1 (Paradis and Schliep, 2019), treeio v. 1.14.0 (Wang et al., 2020)and ggplot2 v. 3.3.3 (Wickham, 2016), tidyverse v.1.3.0 (Wickham et al., 2019), viridis v.0.5.1 (Garnier, 2018), scales v.1.1.1 (Wickham and Seidel, 2020), ggtreeExtra v.0.4.5 (Xu and Yu, 2021), ggstar v.1.0.1 (Xu, 2021), RColorBrewer v.1.1-2 (Neuwirth, 2014), and ggnewscale v.0.4.5 (Campitelli, 2021)packages and Figtree v.1.4.4 (Rambaut, 2009).

### 5. Ploidy level assessment

We assessed the ploidy level of 57 individuals covering the whole phylogeny using flow cytometry following the one-step methodology of Dolezel and Bartos (2005). Briefly, approximately 0.125 cm² of fresh or silica-dried internal standard leaf (*Pisum sativum*’s Ctirad’ 2C= 9.09 pg, Doležel et al., 1998) and 0.125-0.250 cm² of silica-dried fern tissue were co-chopped using a fresh razor blade in a Petri dish containing 1 ml of ice-cold Otto I buffer (0.1 M citric acid, 0,5% Tween 20). The resulting suspension was incubated for 15 min at room temperature (20°C) with occasional shaking and filtered through a 42µm nylon mesh. The filtrate was stained with 1 ml of Otto II buffer (0.4 M Na_2_HPO_4_*12 H_2_0; fluorochrome DAPI: 4 µg/ml) for 1-2 min at room temperature, after which the relative fluorescens was recorded on a CytoFLEX S (Beckman Coulter, Indianapolis, USA). The excitation beam was provided by a laser tuned to a wavelength of 351 nm. Fluorescence emission was detected by a filter permitting passage of light of wavelength of 450 nm. The 2C-values were calculated by comparing the mean *Botrychium* peak to the mean standard peak. Ploidy level were also derived from spore sizes (Popovich et al., 2020) for 14 individuals with genetic proximity to the tetraploid *B. yaaxudakeit* and its maternal donor *B. neolunaria* for which silica-dried material were not available. Spore size of three *B. neolunari*a and five *B. yaaxudakeit* specimens identified by allozymes (Stensvold 2008, D. Farrar unpubl. data) were used as reference. Dried spores were placed in a 1.5 ml Eppendorf tube and suspended in 300 ml of Euparal (Roth, Karlsruhe, Germany) mounting medium. Nine drops of spore solution were equally distributed on a cover slip on the top of which a slide was gently placed. The slides were dried cover slip faced down during a minimum of two weeks before spore measurement. Spores were photographed using inverted microscope (Leica DMI3000B). The spore’s longest length was measured on a minimum of ten spores per individual using LAS v.4.13.Leica Software and ImageJ v.1.52a (Rasband, 1997). Average and standard deviation of spore’s lengths were calculated in R.

### 6. Species delimitation

We considered taxa as independently evolving metapopulations lineages following the concept of de De Queiroz (2007) by applying an integrative approach (Padial et al., 2010). We selected well-supported monophyletic clades to be candidate species (Solís-Lemus et al., 2015; Sukumaran and Knowles, 2017).

Well-supported monophyletic sub-clades nested within species were treated as subspecies when the remainder of the species-level clade did not form coherent units, so that a separation into two or more species was impractical. Species status of clades was further supported by coherent geographical distribution, or distinct ecological features, in combination with previously published information on reproductive isolation between taxa, when available (Wagner and Wagner, 1983b).

### 7. Divergence time

We estimated divergence times within the *B. lunaria* group in a two-step time calibration. First, we estimated the divergence time of the *Botrychium* crown, and second, we inferred the divergence times of the *B. lunaria* group with the *Botrychium* crown age as a secondary derived calibration. In the first step, we analyzed the concatenated alignment of the subfamily Botrychioideae (supplementary File S3, table S3) under strict and uncorrelated lognormal relaxed clock models using BEAST v.1.10.4 (Drummond et al., 2012). The alignment was partition according PartitionFinder 2 output (Table 1) and the site models were unlinked. A birth-death *tree prior* was applied with a given starting tree as *tree model*. The starting tree was built using RAxML-NG and transformed into an ultra-metric tree using the penalized likelihood method (Sanderson, 2002) implemented in the chronopl function of the R ape v. 5.4-1 package. The *clock rate* or the *ucldMean* priors were set with an exponential distribution and defaults parameters. We used two macrofossils to constrain the stem ages of the *Botrypus virginianus* and *Sceptridium* (Bozukov et al., 2010; Rothwell and Stockey, 1989). Morphological analysis related the youngest fossil to *Sceptridium underwoodianum* (Bozukov et al., 2010), whereas the oldest fossil was described as an extinct species, *Botrychium wightonii* Rothwell & Stockey, related to *Botrypus virginianus* (Rothwell and Stockey, 1989). We applied to the *tmrca priors* a uniform distribution with the minimum age of fossils as lower boundaries (56.8 and 20.44 million years, respectively) and the Ophioglossaceae family age estimated by Lehtonen et al. (2017) as upper boundaries (148.9 million years). We set a normal distribution for the *treemodel-rootheight* prior with a mean age of 148.9 million years (Lehtonen et al., 2017) and a standard deviation of 1.0, and we truncated it with an upper boundary in the middle Jurassic (175.6 million years). We constrained the calibrated clades as monophyletic and set *Helminthostachys zeylanica* as outgroup of the subfamily Botrychioideae to facilitate the convergence of the analysis. We ran one analysis per clock model of 20 and 50 million MCMC generations for the strict and the relaxed clock respectively, with parameters sampled every 1,000 generations. Besides, those two analyses were performed without sequence data and without fossil calibrations to control for the influence of the priors on the time-divergence estimates.

Second, we analyzed the A1 alignment containing only unique sequences created by RAxML-NG (supplementary File S4) under strict and uncorrelated lognormal relaxed clock models using BEAST. A yule *tree prior* was applied with a given starting tree as *tree model*. The starting tree was built with the same method than the Botrychioideae starting tree. The *clock rate* or the *ucldMean* priors were set with an exponential distribution and defaults parameters.

We set the distributions parameters using the time-divergence estimates of the genus *Botrychium* given by the analysis of the subfamily Botrychioideae. The *treemodel-rootheight* priors were defined by a normal and a gamma distribution for the strict and the relaxed clock, respectively. The normal distribution had a mean of 14.1 million years, a standard deviation of 1.6 and we truncated it with an upper boundary of 20.7 million years (upper range boundary of *Botrychium* mean). The gamma distribution had a *shape* of 19.5 million years (mean age), a *scale* of 0.7 and we truncated it with an upper boundary of 73.1 million years. We constrained the calibrated clade and *Lunaria* and *Simplex-Campestre* taxon sets as monophyletic to facilitate the convergence of the analysis. We ran one analysis per clock model of 50 million MCMC generations with parameters sampled every 1,000 generations.

The convergence of all analysis was inspected in Tracer v.1.7.1. The effective sample size of every parameter was above 200 and we defined a burn-in of 10 %. We summarized our post burn-in trees using TreeAnnotator v.1.10.4 (Drummond et al., 2012) to generate a maximum clade credibility tree. Besides, we sampled priors for 100 million generations and compared the time divergence estimates with those obtained with the run including the data to ensure the estimates were not driven by priors. Trees were depicted using R ape, phytools v.0.7-70 (Revell, 2012), treeio, ggtree, ggplot2, deeptime v.0.0.5.3. (Gearty, 2021) and strap v.1.4 (Bell and Graeme, 2014) packages. The densitrees were drawn using subsamples of posterior trees obtained with Logcombiner v. 1.10.4 (Drummond et al., 2012).

### 8. Distribution ranges and climatic analysis

We used 511 genotyped individuals with geographical coordinates to define species distribution ranges. The species distribution ranges were visualized using R ggplot2, sf v.0.9-7 (Pebesma, 2018), ggspatial v.1.1.5 (Dunnington, 2021), rnaturalearth v.0.1.0 (South, 2017), and RColorBrewer packages.

We extracted the 19 bioclimatic variables from CHELSA (Karger et al., 2018, 2017) for the collecting localities of a subset of 465 individuals which excluded 46 individuals: 30 individuals with inaccurate geographic coordinates, one individual with uncertain ploidy level, and 15 individuals presumably being introgressed hybrids (Stensvold, 2008; D. Farrar unpubl. data) (supplementary Table S1). The 39 individuals of the *B*. aff. *neolunaria* group were only used to compare the climatic niches of the *B. neolunaria* s.str. and *B*. aff. *neolunaria* groups. The subclade 2 of the *B*. aff. *neolunaria* group had an insufficient number of geo-referenced individuals to be considered in the analysis. Despite of the removal of some accessions, the subset of individuals covered all species and subspecies we recognized. The variable scales varied widely and because variables such as the minimum temperature of the coldest month (BIO 6) had negatives values, log transformation could not be uniformly applied. Thus, we normalized the data using the Z-score standardization. The overall differences between species were visualized for each bioclimatic variable with boxplot representations (supplementary Figure S1). The variables showing differences between taxa were identified both at species and at subspecies level using Analysis of Variance (ANOVA) (supplementary Files S11 to S14). Then, to see which species differed from each other, we performed Tukey post-hoc tests (supplementary File S15). We retained all variables for the analysis at the species level, and 4-16 variables for the analysis at the subspecies level (supplementary Files S16-S19). To visualize the climatic niches of both species and subspecies, we performed principal component analyses (PCA) to account for covariance between climatic variables (supplementary Files S16-S23). The data normalization was done using R *scale* function and analysis were conducted using the R rstatix v.0.6.0 (Kassambara, 2021) and FactoMineR v.2.4 (Lê et al., 2008) packages. The data and results were visualized using the R ggplot2 tidyverse and factoextra v.1.0.7 (Kassambara and Mundt, 2020), cowplot v.1.1.1 (Wilke, 2020), and ggpubr v.0.4.0 (Kassambara, 2020) packages.

Soil pH measurements obtained through field expeditions (see sampling collection section) was refined by measurements realized in the soil biology laboratory of the University of Neuchâtel. Briefly, soil samples were dried at 40°c. Then, one volume of dried soil was mixed with 2.5 volumes of water and stirred 3 times for 5 seconds with 20 minutes between each. The pH was measured 20 minutes after the last agitation using a 914 pH/Conductometer (Metrohm SA, Zofingen, Switzerland). In total, we measured soil pH for 50 populations represented by 54 samples covering eight of the ten clades (supplementary Table S1). The association patterns of soil pH and species were visualized with boxplots using R ggplot2 and tidyverse packages.

## Results

### 1. Locus diversity and plastid-based phylogenetic analysis

The concatenated plastid DNA regions consisted of 2,374 aligned nucleotides of which 4-10 % were parsimony-informative sites (Table 1). A large proportion of individuals (> 80 %) were analyzed at all four target loci (supplementary Table S1). The loci were not equally informative (Table 1). The trnH^GUG^-psbA intergenic spacer and the rpl16 intron were the most variable (114 and 142 parsimony informative sites, respectively), whereas the trnL^UAA^-trnF^GAA^ intergenic spacer was the least informative (81 sites). However, the rpl16 intron presented the lowest sampling completeness due to repeated PCR amplification failure and limited availability of published data (supplementary Table S1, Table 1).

The phylogenetic analysis recovered ten well-supported monophyletic clades inside the *B. lunaria* group (Figure 2, see supplementary Figure S2, S3 and File S5 for tip labels and support values of all branches), consistently arranged into five main groups of clades (Figure 2B, 2C). A first group included three clades (clades 3, 4, and 5), two included each a unique clade (clade 8 and clade 9), another included two clades (clades 2 and 7), and the last included three big clades (clades 1, 6, and 10). The trees resulting from the ML and BI analysis based on the reduced dataset retrieved the same main groups of clades and clades (Figure 3; supplementary Files S6 and S7), except for clades 7 and 2 that were not clustered in the BI analysis, and clade 8 which was only recovered in the ML analysis. However, the relationships between clades were not concordant between the ML and BI analyses (Figure 3).

**Figure 2:**
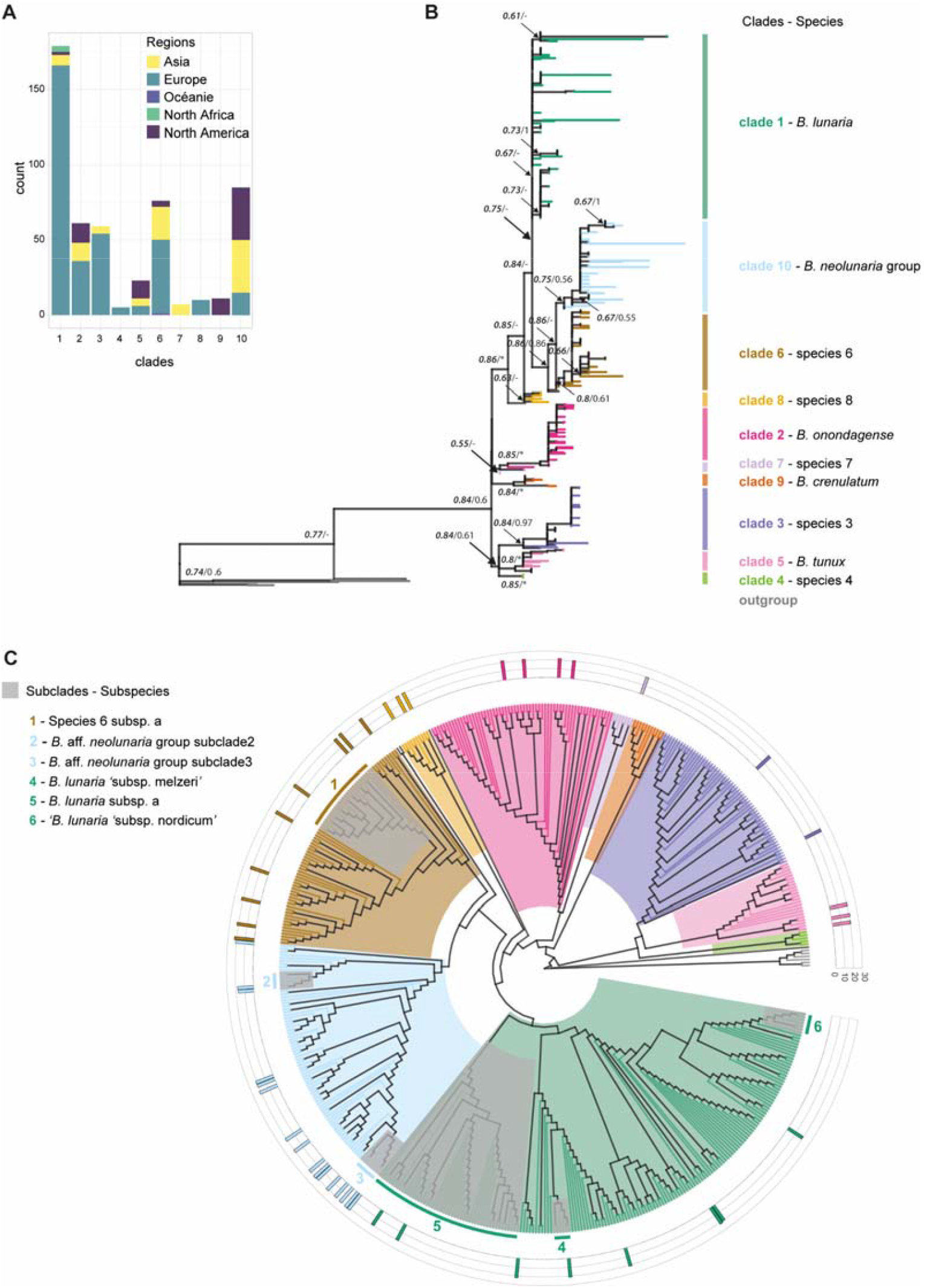
Maximum likelihood (ML) phylogenetic trees depicting the lineages of the *B. lunaria* group with branch support values and genome size distribution. Trees are based on the complete dataset (i.e., including accessions with partial sequence data). Part of the outgroup was pruned of the trees to reduce the root length. (A) Distribution of specimens grouped per clade by large geographic units. (B) Phylogram of the ML tree. ML bootstrap supports (MLBS), and Bayesian inference posterior probabilities (BIPP) are on the left and right, respectively, along the main clade and subclade branches (*i*.*e*., * fully supported, - not applicable; see **supplementary Figure S3, Files S6 and S5** for full BIPP and MLBS values, respectively). Each color represents a distinct clade. (C) Cladogram of the ML tree in fan layout. Branch and background are colored according to clades (tip labels are depicted on **supplementary Figure S2 and S3**). Grey layers on the top of the tree and colored side bars indicate the subclade positions. The layer around the tree shows the absolute genome size distribution given in picograms for the 2C-Values (see supplementary **Table S4** for the values).

**Figure 3:**
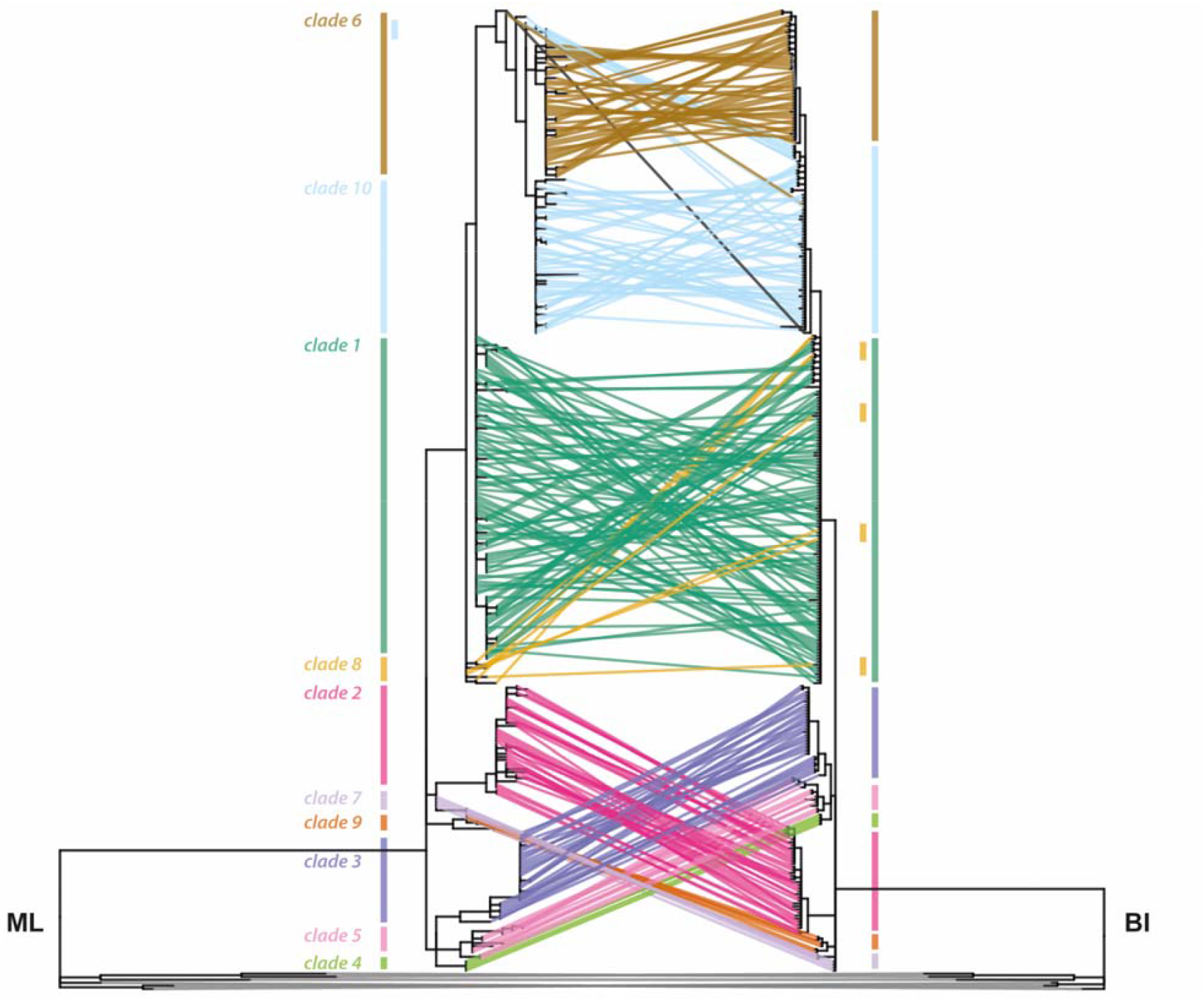
Topological congruence of phylogenetic trees inferred from the reduced dataset using Maximum likelihood (ML) and Bayesian inference (BI) methods. The tree on the right side corresponds to the BI tree and on the left side to the ML tree. Corresponding tips are linked by lines colored according to clade assignment.

The addition of indels slightly increased the branch lengths and the deeper node support in both ML and BI trees (supplementary Figure S4 and Files S8 and S9). The topologies recovered with indels were largely concordant with the analyses based on the reduced trees, and individuals were assigned to the same clades, with few exceptions (supplementary Figure S4). The use of a constrained topology (supplementary File S10) for the ML analysis based on the full dataset allowed the assignment of most individuals with partial sequencing data and low phylogenetic signal. Of the 521 accessions in our study, we were able to reliably assign 509 (97.7 %) to one of the 10 clades. The remaining 12 specimens had uncertain placement, mostly due to incomplete or unreliable sequence data. Eight of these individuals appeared on exceedingly long branches and their placement remained debatable (Figure 2, supplementary Figure S2). Moreover, the assignment of 0ne rogue individual from the Alps (AM-lun33-3-CH) to a clade otherwise restricted to Eastern Asia was suspicious enough to prune it from the final tree.

Some of the ten main clades recovered corresponded to previously recognized species (e.g., *B. tunux* and *B. crenulatum* to clades 5 and 10, respectively), whereas in other cases they combined recognized species, such as *B. neolunaria* and its polyploid derivate *B. yaaxudakeit* (clade 10; Figure 2B, 2C). Some clades showed clear phylogenetic structure within them. We identified six such subclades

(Figure 2C) in the tree resulting from the ML analysis based on the full dataset. Three were within clade 1, namely subclades 4 (*B. lunaria* var. *melzeri*), 5, and 6 (*B. nordicum*), two were in clade 10 (subclades 2 and 3), and one was found in clade 6 (subclade 1). Of these subclades, three are geographically restricted (central Alps, Carpathians, and Baikal; subclades 5, 3 and 2, respectively) and three were also recovered in the BI trees based on the reduced dataset (subclades 4, 2 and 3).

### 2. Ploidy level assessments

The assessment of the ploidy level within the *B. lunaria* group indicated a dominant diploid condition. Absolute genome size showed no evidence of ploidy level variation with 2C-values of 55 specimens between 19.38 and 22.91 pg which attest to a diploid condition (Dauphin et al., 2016; Veselý et al., 2012; Williams and Waller, 2012), except for two individuals from clade 10 subclade 3 with 2C-values of 17.49 and 17.91 (supplementary Table S4). Spore measurements made on 14 additional individuals showed a similar pattern with one exception. Most had length averages between 31 µm and 39 µm (supplementary Table S5) which corresponds to the range found for the diploid *B. neolunaria* (Stensvold and Farrar, 2017). However, one specimen of Caucasus Mountains had a length average of 43 µm which enters the range estimated for the tetraploid *B. yaaxudakeit* (Stensvold et al., 2002).

### 3. Species-clade level classification and candidate species

Well-supported monophyletic clades containing type material or independently identified specimens by allozyme markers (Stensvold, 2008; Stensvold and Farrar, 2017; D. Farrar unpubl. data) of previously recognized species were assigned to the corresponding species, with clades 5, 1, 9 and 2 corresponding to *B. tunux, B. lunaria, B. crenulatum*, and *B. onondagense*, respectively. Among these, *B. onondagense* is currently synonymized with *B. lunaria*. Our species criterion of well-supported monophyly was not applicable to clade 10 because it contained both diploids and polyploids described as distinct species (*i*.*e*., *B. neolunaria* and *B. yaaxudakeit;* Stensvold et al., 2002; Stensvold and Farrar, 2017). In this case, all the specimens of clade 10 were grouped under the name of *B. neolunaria* group, including tetraploids previously identified by allozymes (Stensvold 2002, 2008; D. Farrar unpubl. data) recognized as *B. yaaxudakeit*, specimens previously identified as *B. neolunaria* called *B. neolunaria* s.str. and all remaining specimens known or assumed to be diploids recognized as *B*. aff *neolunaria* group. We did not treat all diploids as the species *B. neolunaria* because they are known to include introgressed hybrids with *B. neolunaria* as maternal parent (Stensvold, 2008; Stensvold and Farrar, 2017). Thus, while they cluster in the same cade, they have different genetic backgrounds and very variable morphology (V. Mossion unpubl. data). The remaining five clades are treated as unnamed candidate species and named by their clade numbers (species 3, 4, 6, 7 and 8). We further recognized four monophyletic clades nested within species that are treated here at subspecies level for consistency, including the taxa recognized as species level as *B. nordicum* and as variety level as *B. lunaria* var. *melzeri* (Stensvold and Farrar, 2017). We thus use the names *B. lunaria* “subsp. *nordicum*” and *B. lunaria* “subsp. *melzeri*”, but without formal recombination. The subclades nested within the *B. neolunaria* group (clade 10) are not treated at the subspecies level as their species assignment is unclear.

### 4. Divergence time estimates

Our age estimates suggest that the radiation of the *B. lunaria* group started at the end of the Miocene and extended to the Pliocene, about 45 million years (myr) after the genus *Botrychium* diverged from its sister genera *Japanobotrychium* and *Sceptridium* (Figure 4A, supplementary Figure S5). The results are comparable between the two clock models even though the uncorrelated lognormal relaxed clock model recovered higher divergence time estimates for both genus- and species-level analyses (Figure 4B, supplementary Figures S5, S6, and S7). For example, the mean estimates of the crown ages of the genus *Botrychium* were 19.5 (95 % HDP 7.1-22.6 myr) and 14.1 (95 % HDP 8.7-20.8 myr) under the uncorrelated lognormal relaxed clock model and the strict clock model, respectively. The *B. lunaria* group arose in the Pliocene epoch (mean 3.6 myr, 95 % HDP 2.38-5.053 myr) while the species within mostly diverged from one another during the Gelasian, the first Pleistocene age at 2.58-1.8 myr (Figure 4C, supplementary Figure S6, and S7). The divergence times of the subspecies were estimated within the 95 % interval confidence only for subspecies “a” of species 6 and for *B. lunaria “*subsp. *nordicum*” due to missing data, rogue individual, and a lack of a sufficient number of unique sequences. They diverged from the remaining of the clade at the end of the Calabrian and Chibanian ages, the second and third quarter of the Pleistocene epoch at ∼1 myr and ∼0.5 myr, respectively (Figure 4C, supplementary Figure S6 and S7).

**Figure 4:**
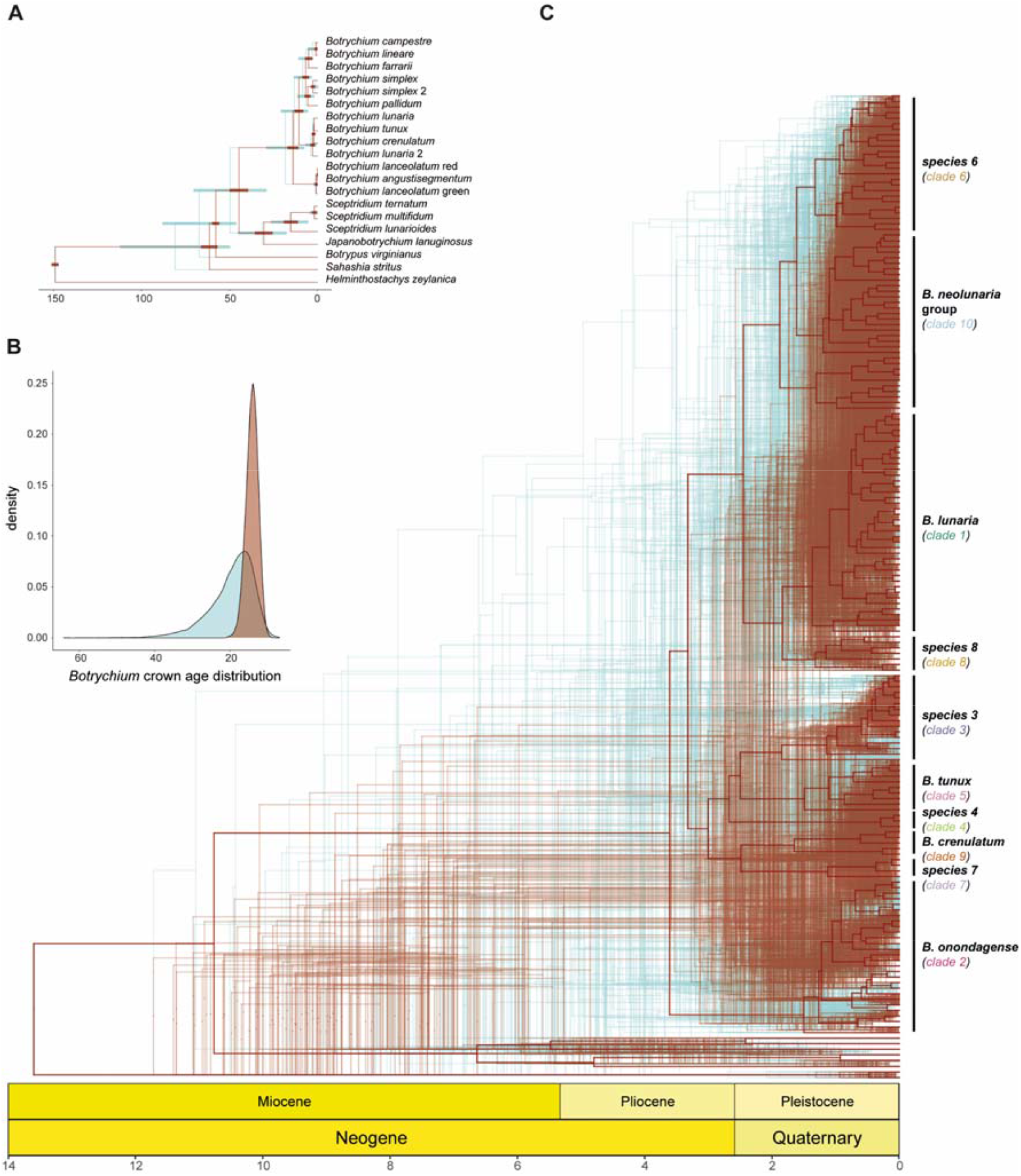
Divergence time estimates of the subfamily Botrychioideae and of the species of *B. lunaria* complex. Time scales are in million-year unit. The red color refers to the divergence time analysis ran under the strict clock model, and the blue color to the relaxed clock model. (A) Time calibrated phylogenies of the subfamily Botrychioideae. The 0.95 Highest Posterior Density (HPD) is represented by bars at the calibrated nodes. (B) Distribution of the *Botrychium* crown age density estimated by the Botrychioideae time-divergence analysis. (C) Time calibrated phylogeny of the *B. lunaria* complex. The uncertainty around the node ages is displayed by density trees. The density trees were each drawn on subset of 90 posterior trees. The geological time scale shows the periods and the epochs and follows the International Chronostratigraphic Chart (2018; https://stratigraphy.org).

### 5. Distribution ranges of species in the B. lunaria group

Our classification into species results in the recognition of seven species in North America (USA, Canada, Greenland), seven in Asia, one each in northwestern Africa and Oceania, and eight to nine (considering the potential tetraploid from Georgia) species in Europe (Figure 2A, Figure 5A-C). Of the 11 candidate species found among our samples, five (species 4, 7, 8, *B. crenulatum*, and *B. yaaxudakeit*) had narrow geographical ranges. These species were restricted to northern and eastern Europe, the Himalayas, as well as western and northern North America. The remaining species had wider distributions. For instance, *B. lunaria* (Figure 5A) was recovered from northwestern Africa to Scandinavia and Central Asia. *Botrychium tunux* and *B. onondagense* both had a disjunct distribution in Europe and North America with a disjunction between Eastern European and Central Asian populations, and no sample from western North America was assigned to *B. onondagense*. Species 3 and 6 were both found from Central and Eastern Europe to eastern Asia with a marked disjunction in Central Asia for species 3 and a few records in Alaska for species 6. The most widespread species-level group was the *B. neolunaria* group (Figure 5B), which we recorded across almost the entire range of the *B. lunaria* group.

**Figure 5:**
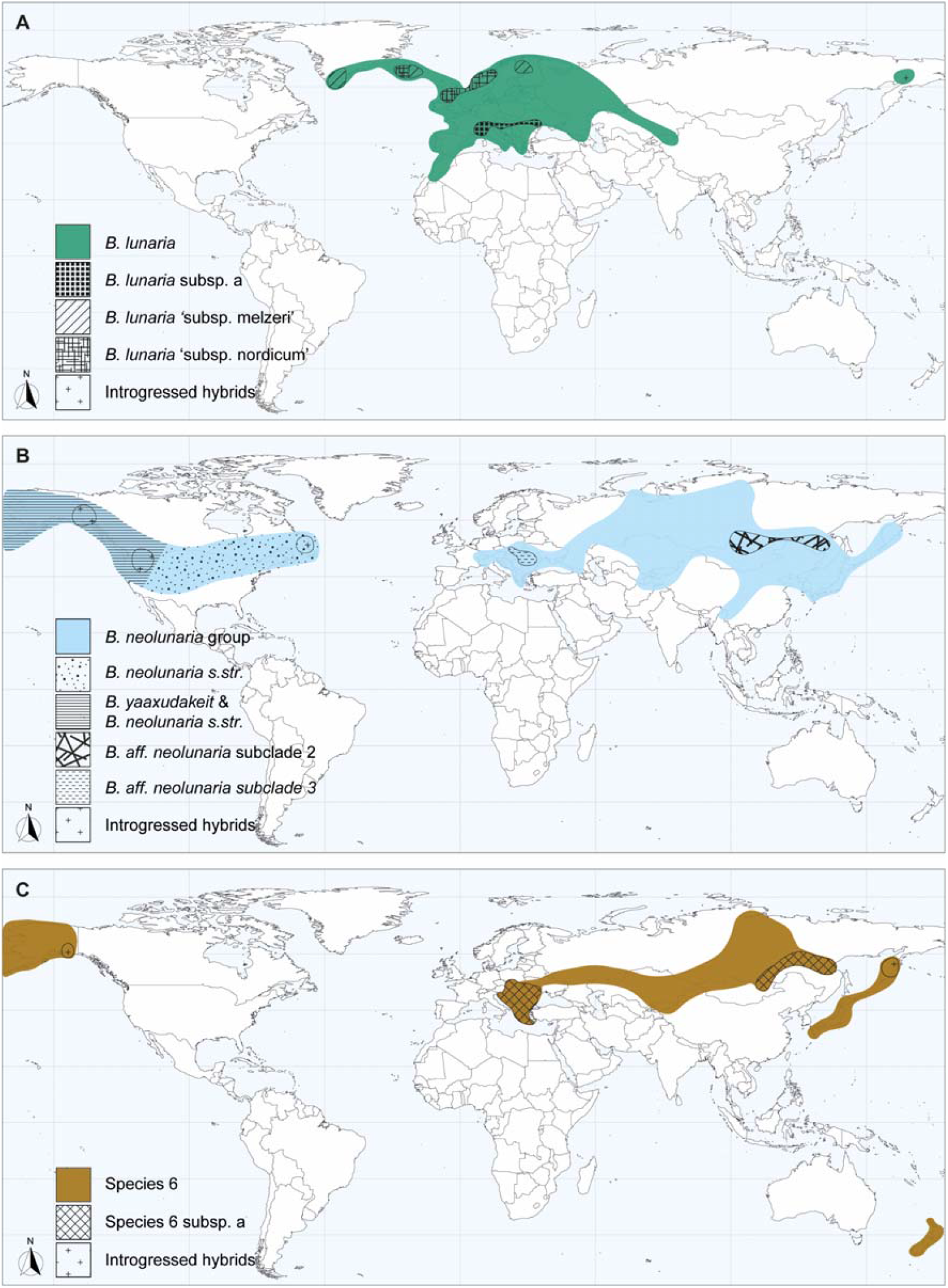
Geographic distributions of the species of the B. lunaria group. (A) Global distribution of B. *lunaria* (clade 1). (B) Global distribution of the *B. neolunaria* group (clade 10). (C) Global distribution of the species 6 (clade 6).

### 6. Climatic and soil analyses

The climatic niches largely overlapped between taxa, albeit with slight local differences. The principal component analysis (PCA) clustered the species into two groups. The first group consisted of species 1, 3, 8, and *B. crenulatum*, and the second group of species 2, 4, 6, 7, *B. neolunaria* s.str., and *B. tunux* (Figure 6). The first component (PC1, 40.0 %) separated the species by temperature-related variables (BIO 1, 11, and 6) showing a gradient of strength of winter from mild winters (first group) to strong winters (second group) (Figure 6A-B). The second component (PC2, 23.8 %) was correlated with precipitation-related variables (BIO 12, 13, 14, 16, 17, 19) and temperature seasonality (BIO 7 and 4), corresponding to a gradient of climatic continentality from less seasonal climates with high annual precipitations in areas close to oceans to strongly seasonal climates with low annual precipitations in continental regions. Within both groups, species were arranged along this continentality gradient. For example, in the second group species were arranged from species 7 (oceanic) to species 4 (continental) (Figure 6C-D). The third component (PC3, 12.6 %) was related to temperature-related variables (BIO 10, 5 and 1) expressing a gradient of summer temperature, with the first group under cooler summer climates than the second group. PC3 was also correlated to diurnal temperature fluctuations and their constancy through the year (BIO 2 and 3; supplementary Figure S8). Thus, the PCA recovered that *B. crenulatum* and *s*pecies *7* occur in habitats with a large range of diurnal and annual temperature fluctuations, with the former subjected to even stronger temperature variations.

**Figure 6:**
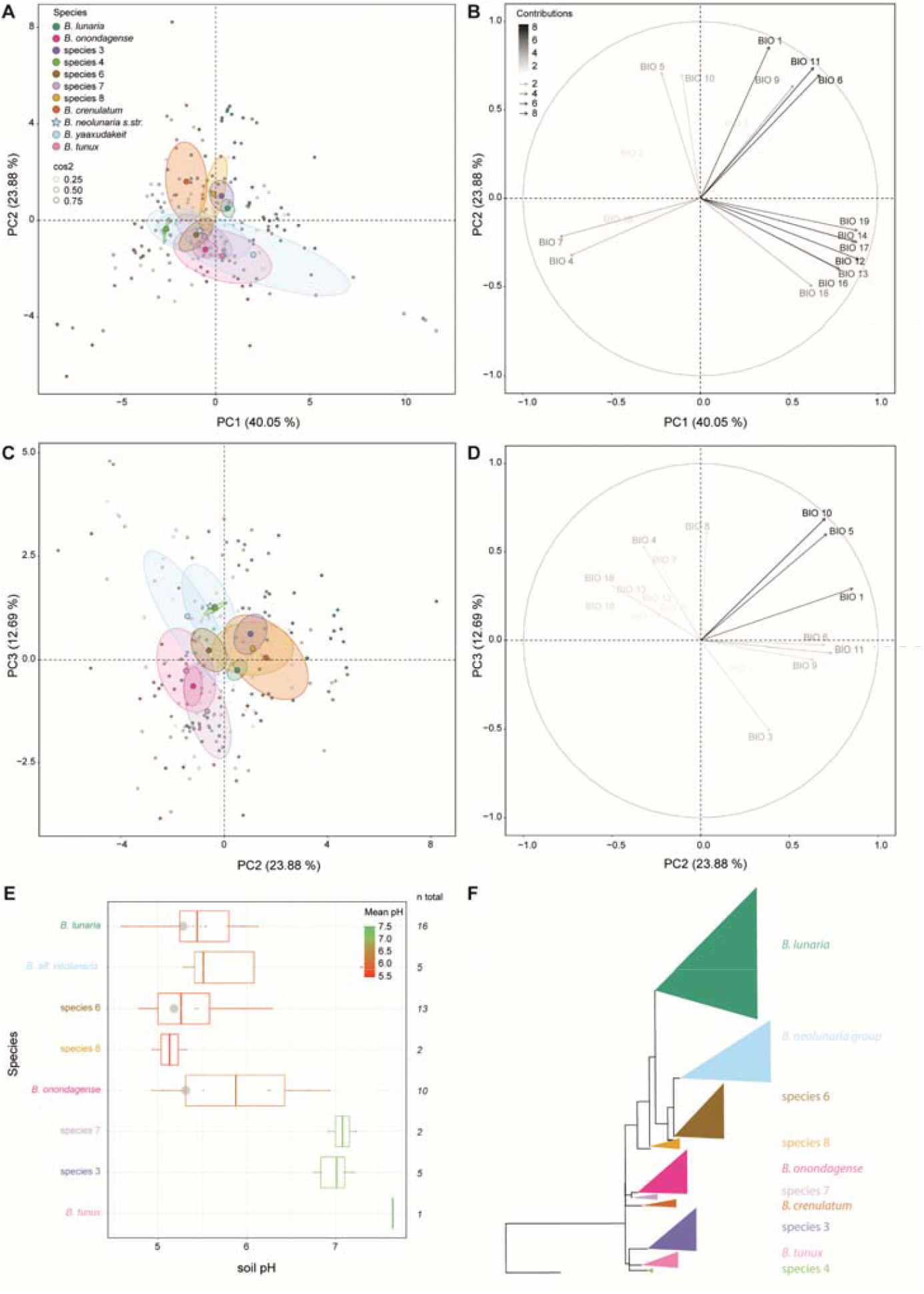
Ecological differentiation between species of the *B. lunaria* group. (A-D) Principal Component Analyses (PCA) of climatic factors comparing the species occurrences. Point transparencies increase with lower cos2 values. Points and 95 % confidence ellipses are colored according to clade assignment, with centroids indicated by larger circles. Arrow transparency represents the variable contributions to the PC. The definitions of the CHELSA bioclimatic variables are given in the **supplementary Table S6**. (A) Individual factor map showing the PC 1 and 2. (B) Variable factor map showing the PC1 and PC2. (C) Individual factor map showing the PC 2 and 3. (D) Variable factor map showing the PC2 an PC3. (E) Boxplots of soil pH. The boxplot colors represent the pH mean per species. The grey dots show the distribution of the individual pH values, and their size is relative to the number of individuals with similar pH values. The sample size per species is given by ***n***. (F) ML phylogenetic tree depicted with collapsed clades. Triangle height relates the clade sizes, and the triangle width the maximum branch length

We also observed differences in the climatic niches between subspecies (supplementary Figure S9) and between the *B. neolunaria* s.str. and *B*. aff. *neolunaria* groups. The PCA within species 1 showed substantial climatic niche differentiation explained by seasonality (PC1, 40.3 %), high temperatures (PC2, 26.8%), and the strength of the winter (PC3, 15.7 %). In brief, *B. lunaria* “subsp. *melzeri*” occurred in dry climates with a strong seasonality and cold winters, *B. lunaria “*subsp. *nordicum”* in wet climates with low seasonality and low annual temperatures, and the two others under more temperate climates. The PCA within species 6 revealed similar importance of the seasonality (PC1, 62.3 %) and precipitation (PC2, 16.9 %), with the nominal subspecies in dry habitats characterized by medium seasonality and cold winters, and subspecies 1 in wet habitats characterized by low seasonality and mild winters. The PCA of the B. *neolunaria* showed differences between *B. neolunaria* s.str., and *B*. aff. *neolunaria* group related to summer temperature (PC1, 50.78 %), and seasonality (PC2, 26.2 %) with *B. neolunaria* s.str. and the *B*. aff. *neolunaria* group in climate with hot and dry summer, and higher summer temperatures for the former, and except subclade 3 individuals in climates with cold and wet summers.

Soil pH showed differences between species even though the sample size was limited. *Botrychium lunaria*, species 6, and species 8 were only found on acidic soils, whereas *B. tunux*, species 3 and species 7 were recovered exclusively from neutral soils. *Botrychium* aff. *neolunaria* group was mostly found on both acidic and neutral soils (Figure 6E-F). Interestingly *B. onondagense* was found across the whole range of pH values recorded in the *B. lunaria* group with a trend towards the acidic soils.

## Discussion

Based on over 500 sampled individuals, we provide the first global molecular phylogenetic analysis of the *Botrychium lunaria* group. The lineage diversity recovered is congruent with previous, geographically more limited studies and uncovered three novel monophyletic clades, resulting in the recognition of 11 candidate species and 4 candidate subspecies. These are evenly distributed among whole northern hemisphere, with seven or eight species per continent. We found little evidence that polyploidization played a role in the diversification of the group, although hybridization appears to be important in one clade. The time-calibrated phylogeny shows a concordance of Pleistocene climatic oscillations with the radiation of the group. Moreover, climate and soil pH showed slight differences among candidate species and subspecies, suggesting that ecological drivers played a role in the diversification of the *B. lunaria* group.

We freely acknowledge that one limitation of our study is that it only includes non-coding plastid regions, so that hybrids and polyploid individuals may not have been recognized in the absence of nuclear markers. However, as discussed below, the clades recovered by us make up consistent geographical and ecological units, lending support to the notion that they are real evolutionary entities.

### Taxonomic diversity in the B. lunaria group

Our plastid-based phylogeny recovered ten well-supported clades, of which three are novel (clades 6, 7, and 8) compared to earlier studies based on the same non-coding plastid regions (Dauphin et al., 2017, 2014; Maccagni et al., 2017). The remaining seven clades are consistent with previous studies even though their relationships differ. Clades are arranged in five main groups in our analysis against two in the previous studies. For example, clades 9 (*B. crenulatum*) and 2 were recovered separated from each other, whereas they were previously retrieved as sister clades (Dauphin et al., 2017).

The ten clades found in the *B. lunaria* group include 17 candidate taxa. Eight of these correspond to currently accepted taxa, namely *B. lunaria* (Swartz, 1801), *B. crenulatum* (Wagner and Wagner, 1981), *B. tunux* (Stensvold et al., 2002), *B. yaaxudakeit* (Stensvold et al., 2002), *B. nordicum* (Stensvold and Farrar, 2017); our *B. lunaria* “subsp. *nordicum*”), *B. neolunaria* (Stensvold and Farrar, 2017), and *B. lunaria* var. *melzeri* (Stensvold and Farrar, 2017); our *B. lunaria* “subsp. *melzeri*”). Four other clades have previously been recovered in phylogenetic analyses but have not been described or were synonymized, namely *B. onondagense* (Underwood, 1903) (Clade 2), synonymized with *B. lunaria* by Clute (1905) as well as species 4, species 3, and *B. lunaria* subsp. “a” (Dauphin et al., 2017, 2014; Maccagni et al., 2017). Finally, four clades are novel candidate taxa (species 6, 7, 8, species 6 subsp. “a”). This increased number of taxa is the result of our geographically more extensive sampling and overall increased sample size.

The consistency of the ten species-level clades in the ML and Bayesian analyses, despite different arrangements among the clades in the two analyses, in combination with their ecological, geographical, and morphological distinctness (Stensvold et al., 2002; Stensvold and Farrar, 2017; Wagner and Wagner, 1981; V. Mossion unpublish. data), supports the notion that these clades are best treated at species level. Although hybrids have been documented in the *B. lunaria* group (Stensvold, 2008), the fact that co-occurring species such as *B. neolunaria* and *B. tunux* in North America (Figure 1Ag) conserve their morphological distinctiveness suggest these species are reproductively largely isolated (Wagner and Wagner, 1983b). Unfortunately, directly testing for reproductive isolation between lineages of the *B. lunaria* group by breeding experiments is technically restrained by the difficulty of cultivating the subterranean gametophytes (Campbell, 1911; Whittier, 1972, 1981; Mossion unpubl. data). We are nevertheless confident that our treatment of the ten major clades recovered in our analyses as species is sound, along with the recognition of allotetraploid *B. yaaxudakeit* as an eleventh species.

The taxonomic treatment of clades nested within the species-level clades is less straightforward, as exemplified by the case of *B. nordicum*, here referred as *B. lunaria “*subsp. *nordicum”*, which was found to be deeply nested within *B. lunaria*, corroborating previous findings (Dauphin et al., 2017). This taxon was described at species level based on morphological distinctness and unique alleles separating it from *B. “lunaria”* (Stensvold and Farrar, 2017). In our analysis, this taxon was recovered as a monophyletic clade nested within *B. lunaria* so that treating it as specifically distinct would render the remainder of *B. lunaria* paraphyletic. Our divergence time estimates revealed that *B. lunaria “*subsp. *nordicum”* has only recently derived from the rest of *B. lunaria*, and its nested placement is thus likely the result of incomplete lineage sorting (Pamilo and Nei, 1988; Wendel and Doyle, 1998). For consistency, we treat *B. lunaria “*subsp. *nordicum”* at subspecies level, but considering its morphological distinctness and the allozyme patterns (Stensvold and Farrar, 2017), further investigations involving nuclear sequencing data are necessary to decide on any taxonomic rank modification. The same is true for the other subspecies recognized here.

Finally, the *B. neolunaria* group presents special taxonomic problems. Not only does it include allotetraploid *B. yaaxudakeit*, which shares the plastid genome with it maternal parent *B. neolunaria* and thus cannot be distinguished with our data, but there also appears to be considerable genome size variation within this group. Thus, our genome size measurements for Eurasian individuals of the *B. neolunaria* group showed much lower 2C-values (17.49 – 21.88 pg) than reported for “true” *B. neolunaria* from North America (27.26-27.75 pg; (Dauphin et al., 2016). To date, no record of “true” *B. neolunaria* identified by allozyme markers is known from Eurasia, even though hundreds of plants have been analyzed (Stensvold, 2008). However, a considerable amount of fertile introgressed hybrids between *B. neolunaria* and B. “*lunaria”* has been recorded from Eurasia. We propose that the differences in genome size in the *B. neolunaria* group reflect the presence of several taxa originating from multiple hybridization events between *B. neolunaria* and one or several other species from the *B. lunaria* group. We would not be surprised if this group contains several species-level taxa, but our plastid markers only indicate the maternal donor, and do not allow to test this hypothesis. Further studies involving nuclear markers are needed to address the evolutionary history and taxonomic diversity of *B. neolunaria* group.

### Global geographical distribution of the B. lunaria group

Based on intensive sampling in Western and Central Europe, Maccagni *et al*. (2017) proposed that continental Europe is the center of diversity of the *B. lunaria* group. However, our geographically more exhaustive sampling shows that species numbers are quite evenly distributed between continents, with seven species in North America, eight in Europe, and seven or eight in Asia. Importantly, our sampling is less dense in Asia (94 versus 346 specimens in Europe), with large areas yet to be sampled. It is thus conceivable that further species diversity might occur in Asia.

Dauphin et al. (2017) hypothesized that the *B. lunaria* group may have originated in Asia, since a clade mainly composed of Asian specimens was sister to the remainder of the group. Testing this hypothesis requires to firmly resolve the relationships between clades, which is not the case in our analysis. Indeed, the relationships between clades were inconsistent between the maximum likelihood (ML) and Bayesian inference (BI) methods. For example, clade 7, exclusively composed of Asian specimens, was sister to all the others in our BI trees, whereas the group of clades 4, 5 (*B. tunux*), and 3, which contain mostly European and North American specimens, was recovered as such in the ML trees. It appears that our increased sampling size is not sufficient to resolve the deeper nodes and that additional sequencing data, preferably including nuclear data, are needed to address question on the geographical origin of the *B. lunaria* group.

Considering the intensive previous studies in North America (Stensvold, 2008; Stensvold et al., 2002; Stensvold and Farrar, 2017; Wagner and Wagner, 1981), unsurprisingly most of the newly found candidate species have ranges restricted to Europe or Asia. Our study extends the known geographic distribution of most widespread species but confirms the narrow distributions of *B. crenulatum* and *B. lunaria* subspecies “a” (Dauphin et al., 2017; Stensvold, 2008; Wagner and Wagner, 1981). Perhaps the most surprising range extension concerns the *B. neolunaria* group, which was previously only known from Eastern Asia, Oceania, and North America (Dauphin et al., 2017; Stensvold and Farrar, 2017) but was recorded by us across all Eurasia. However, disentangling the distribution of introgressed hybrids and “true” *B. neolunaria* is not possible here as both share the same plastid genome. The same statement applies for the distribution of *B. yaaxudakeit*, the derivative polyploid of *B. neolunaria*. However, genome size estimates showed no evidence of tetraploid plants that might be referable to *B. yaaxudakeit* in Eurasia, except for a single specimen with large spores from Georgia, whose taxonomic identity remains unresolved.

### Limited roles of polyploidization and hybridization in the diversification of the B. lunaria group

A previous study of other clades in the genus *Botrychium* has uncovered complex polyploid networks, revealing an important role of allopolyploidization in species formation in the genus (Dauphin et al., 2018). In contrast, we found little evidence of ploidy variation within the *B. lunaria* group despite extensive sampling and additionally selecting morphologically aberrant specimens that might be hybrids. In combination with previous studies (Dauphin et al., 2016; Veselý et al., 2012; Williams and Waller, 2012), we thus show that the diploid condition is predominant in the *B. lunaria* group. This finding indicates that ploidy variation is not a major driver of diversification in the *B. lunaria* group, with the exception of allotetraploid *B. yaaxudakeit* (Stensvold et al., 2002). Nevertheless, as detailed above, hybridization may have led to still poorly understood diversification in the *B. neolunaria* group.

### Pleistocene climatic oscillations influenced the radiation of the B. lunaria group

Our study shows that the current species diversity of the *B. lunaria* group originated during the last 3 million years, suggesting a possible link with the massive Pleistocene climatic fluctuations. Large parts of the current ranges of the species in the *B. lunaria* group were covered by ice during glacial times, including the entire known range of species 4 in boreal Sweden, and most of the habitat of the species occurring in the Alps (Ehlers et al., 2018; Ivy-Ochs, 2015). This is a well-known situation for alpine and boreal plant species which has frequently been invoked as driving the divergence of specific and intraspecific genetic differentiation by repeated cycles of extinction, migration, allopatry, and sympatry (Tribsch and Schönswetter, 2003; Westergaard et al., 2011). In the *B. lunaria* group, this has not resulted in the formation of primarily allopatric species, as for example in the European gentians (Hungerer and Kadereit, 1998), but rather in the formation of numerous widespread species that mostly broadly overlap geographically and that often co-occur. For example, in the Swiss Alps, *B. tunux* has mostly been found in mixed populations with species 3, while *B. lunaria* subsp. “a” and species 3 were also recovered in sympatry (Maccagni et al., 2017). In our study, we found similar patterns, with for example *B. onondagense* and species 3 co-occurring at a site in Romania, or species 6 and *B. lunaria* co-occurring at several sites in Bosnia and Herzegovina. This co-occurrence of species is certainly a result of the high dispersal ability of ferns by their spores (Barrington, 1993), especially long-distance dispersal by wind, and attests to a highly dynamic history of population movements which cannot be reconstruct based on plastid data. Although hybrids are known to occur between species of the *B. lunaria* group (Stensvold, 2008), it is evident that species identity is not primarily maintained by limited gene flow between geographically remote populations, but rather by pre- or postzygotic mating barriers which further supports the treatment of clades as species (Wagner et al., 1985). However, whether original species divergence took place in sympatry or parapatry, or whether it was kick-started by allopatry, remains open to speculation.

### Climatic and edaphic differentiation between species

To our knowledge, no study has directly investigated the importance of ecological factors as drivers of the diversification in the *B. lunaria* group. However, Maccagni et al. (2017) observed possible habitat differences between two species (*B. lunaria* and our species 3) suggesting a role of ecological segregation. Also, the distribution of arbuscular mycorrhizal fungi (AMF) on which the growth of *Botrychium* gametophytes relies, is driven by abiotic factors (Davison et al., 2021; Sandoz et al., 2020) which implies potential indirect edaphic preferences.

We found that the climatic niches of both species and subspecies within the *B. lunaria* group showed a certain degree of differentiation, but with marked overlap between many species. The main differentiation occurred along gradients of elevation (as reflected by various temperature-related variables) and climatic continentality. The two main groups of species recovered in the PCA on climatic factors did not correspond to the main group of clades recovered in the phylogenetic analyses, showing that low- and high-elevation species occur in several clades and that their differentiation was not driven by elevational specialization. Unfortunately, the high uncertainty regarding species-level relationships within the main groups precludes any conclusions about evolutionary tendencies related to species diversification within these clades. In any case, the conclusion from these analyses would be that while there is some climatic niche differentiation between species, this is relatively weak and unlikely to suggest that climatic divergence would be a major driver of diversification in the *B. lunaria* group.

Interestingly, however, in the analyses of climatic niches within variable species, we found that subspecies were mostly better segregated in climatic niche space than the species. This likely reflects the fact that analyses within single species encompass a narrower range of climatic conditions as well as fewer taxa, allowing for clearer segregation between taxa. Yet, considering that differentiation at subspecies level might be a precursor to species-level differentiation, and contrary to our conclusions above, this would suggest that climatic niche differentiation might indeed play an important role in the diversification of the *B. lunaria* group.

Our soil pH data from a subsample of sites also reveal patterns of edaphic preferences between species, with some species recorded only on neutral soils (*B. tunux*, species 3, species 7) and others on acidic soils (*B. lunaria*, species 6, species 8), although *B. onondagense* and *B*. aff. *neolunaria* group occurred across the entire pH range. This suggests that there might be some degree of edaphic differentiation between species. In North America, *Botrychium* species have been found associated with calcareous bedrock, basalt bedrock and coastal soils influenced by ocean spray (D. Farrar pers. obs.) with a higher species abundance and richness on neutral or basic soils (Farrar et al., 2017). The soil pH data from North America were extrapolated from bedrock substrates, but our data showed partial dissociation between bedrocks and soil pH. Indeed, two third of the populations with acidic soil pH were found on limestone, dolomite, marble, sandstone, or conglomerate bedrocks, suggesting small scale soil heterogeneity (Hartmann and Moosdorf, 2012).

Soil pH and temperature have been shown to explain the realized niche of arbuscular mycorrhizal fungi (AMF) (Davison et al., 2021), with clear differences among *Glomus* species, which are known to be the main group of AMF associated with *Botrychium* species (Sandoz et al., 2020; Winther and Friedman, 2007). In the Alps, AMF communities exhibit distinct composition in acidic and neutral soils (Sandoz et al., 2020). As the growth of *Botrychium* is dependent on their AMF symbiosis, it might well be that pH preferences between species reflect association with distinct Glomeraceae species and that the co-evolution between AMF and *Botrychium* played a role in the diversification of the *B. lunaria* group.

In conclusion, we find that there is some ecological differentiation between species of the *B. lunaria* group, but evidence of this is rather weak and at present we are unable to identify it as a prime driver of speciation in the group. Even though limited, the soil data available to us evokes potential small-scale ecological differentiation that may not be recovered by our climatic data (at 1 km^2^ resolution). We suggest that in-depth studies of ecological conditions at sites where several species co-occur might help to further understand the role of ecological specialization in this group.

## Conclusions

Based on a global sample of over 500 specimens, our study reshapes the understanding of the species-level diversity within the *B. lunaria* group and the geographical distribution of its diversity. We find that the timing of diversification corresponds to that of Pleistocene climatic oscillations, suggesting that diversification was mainly driven by repeated cycles of dispersal and extinction. This may then have been combined with some cases of polyploidization (*B. yaaxudakeit*, perhaps a further one in Georgia) and hybridization (in the *B. neolunaria* group), as well as slight climatic and edaphic differentiation between many species and subspecies. Although most of species-level taxa we recognize are 2-3 million years old, the fact that we also found several much younger, genetically, and climatically distinct sub-clades, which we treat at subspecies level, suggests that diversification in the group is ongoing. Future studies should focus on clarifying the clade relationships within the *B. lunaria* group, on understanding the evolution and taxonomic implications of variation within the *B. neolunaria* group, and on documenting the role of small-scale ecological differentiation between species.

## Acknowledgements

For their help organising and carrying out field work, we are in debt to Alexandru Coltoiu and the Bucegi Natural Park, Calimani National Park administration (Romania), Claudia Danau and the Retezat National Park (Romania), Clément Duckert, Cun-Feng Zhao, Daniel Ston, Kenan Ćatović, Kuban Uulu Zholdoshbek and the administration of Sarychat-Ertash State Nature Reserve (Kyrgyzstan), Luca Lässer, Marine Ramirez, Quentin Dubois, Sabin Neatu and the Ceahlau National Park (Romania), Sandra Grünig, Sergei Aleksandrov and the Central Balkan National Park (Bulgaria), Shamil Shetekauri, Siqi Liang, Tolkha Shetekauri and the team of Tbilisi Botanical Garden (Georgia), Xianchun Zhang, and Zikriyo. We are also thankful to Ahmed Ouhammou (MARK), Alessio Maccagni, Alexey P. Seregin (MW), Andreas Beiser, Dauphin Benjamin, Iowa State University (ISC), Filippo Prosser, Hedwig Meindl, Isabelle Chanaron (AIX), Markus Grabher, Natalia Gamova, Robin Walls, Sigitas Juzėnas, Siri Birkeland, Teddy Dolstra, Vincenzo Ferri, Wittmann Helmunt, Xian-Yun Mu, and Xianchun Zhang (PE) for providing specimens. Many thanks to Amandine Pillonel, Ophélie Gning, Pierre-Emmanuel DuPasquier, Luyinda Lukau, Bondo Mateus, and Benite Abayo who helped with the molecular laboratory work. Amandine Pillonel prepared the soil samples and made the ex-situ pH measurements. Elke Kessler conducted the flow cytometry analyses. We thank Pierre-Emmanuel DuPasquier, Libing Zhang, and Liang Zhang for the advanced access to Botrychioideae sequencing data. We also acknowledge Sabina Moser Tralamazza and Ursula Oggenfuss for their thoughtful revisions of the manuscript.

## Funding

This work was supported by grants from the ‘Fond des donations’ and the ‘Subvention égalité’ of the University of Neuchâtel.

## Abbreviations

MARK: Cadi Ayyad University herbarium
MW: Moscow State University herbarium
ISC: Iowa State University herbarium
AIX: Muséum d’Histoire Naturelle d’Aix-en-Provence herbarium
PE: Institute of Botany, Chinese Academy of Sciences herbarium
NEU: University of Neuchâtel herbarium
Z/ZT: herbarium of the University of Zurich and the Federal Institute of Technology in Zurich

## Competing interests

none

## Author contributions

VM, JG and MK designed the study. JG, MK and VM acquired the funding. VM organized and performed the fieldwork. DF provided essential type and identified reference plant material. EK, VM, DC and MK performed the analysis. MK, VM and DF interpretated the results. VM and MK wrote the first draft with substantial inputs from DC and DF. All co-authors reviewed the manuscript and approved the final version.

## Data accessibility

Sequences generated in this study were deposited on Genbank. The accession numbers are given in the Supplementary Table S1 and S3. Alignments, and phylogenetic trees are available as supplementary Files S1-S10 (https://doi.org/10.5281/zenodo.6531504).

## Notes

### Competing Interest Statement

The authors have declared no competing interest.

