## Supplementary information for "Global diversification of the common moonwort ferns (*Botrychium lunaria* group, Ophioglossaceae) was mainly driven by Pleistocene climatic shifts"

Running title: *Botrychium lunaria* diversification

Vinciane Mossion^1^, Erik Koenen^2^, Jason Grant^1^, Daniel Croll^1^, Donald R. Farrar^3^, Michael Kessler^2^

^1^ Laboratory of Evolutionary Genetics, University of Neuchâtel, Neuchâtel, Switzerland

^2^ Department of Systematic and Evolutionary Botany, University of Zürich, Zurich, Switzerland

^3^ Department of Ecology, Evolution and Organismal Biology, Iowa State University of Science and Technology, United-States of America

*Supplementary Figures*

[Available from Zenodo: https://doi.org/10.5281/zenodo.6531504]

**Supplementary Figure S1: Boxplots of the differences between species for each bioclimatic variable**.

**Supplementary Figure S2: Phylogram of** **the Maximum likelihood (ML) tree including all accessions with tip labels**. The tree is depicted in the rectangular layout. Tip labels correspond to the sequence code, type information (*i.e.,* taxa name and type status) when applicable, the region of origin and the three letters ISO code of the country. For further information on accessions refer to the supplementary Table 1. The tip colors indicate clade assignment, and the Bootstrap supports of the main clades and subclades are specified on left node sides.

**Supplementary Figure S3: Posterior probabilities (PP) of** **the Bayesian (BI) tree based on reduced dataset.** Phylogram of the BI tree in rectangular layout. The PP are depicted on the right side of nodes. Tip labels correspond to the sequence code, type information (*i.e.,* taxa name and type status) when applicable, the region of origin and the three letters ISO code of the country.

**Supplementary Figure S4: Effect of indels on the trees’ topology inferred from the reduced dataset.** (A) Tanglegram of the trees inferred using Maximum likelihood (ML) method. The tree on the right side corresponds to the ML tree inferred with indels and on the left side to the ML tree inferred without indels. Corresponding tips are link by lines colored according to clade assignments. (B) Tanglegram of the trees inferred using Bayesian (BI) method. The tree on the right side corresponds to the BI tree inferred with indels and on the left side to the BI tree inferred without indels. Corresponding tips are link by lines colored according to clade assignments.

**Supplementary Figure S5: Time calibrated phylogenies of the Botrychioidae sub-family genus.** On the left-side time divergence analysis ran under strict clock model and on the right-side time divergence analysis ran under relaxed clock model. The dark red and cadet blue bars represent the height of 0.95 HPD. The height medians are specified above the bars. The black dots indicate a posterior probability above 0.95. The time scales are in million-year unit.

**Supplementary Figure S6: Time calibrated phylogeny of the *B. lunaria* complex inferred under the strict clock model.** The dark red bars represent the height of 0.95 Highest Posterior Density (HPD) of the calibrated nodes. The height medians are specified above the bars. The black dots indicate a posterior probability equal or above 0.95 and the grey dots show a posterior probability comprises between 0.90 and 0.95. The time scale is in million-year unit.

**Supplementary Figure S7: Time calibrated phylogeny of the *B. lunaria* complex inferred under the relaxed clock model.** The cadet blue bars represent the height of 0.95 HPD of the calibrated nodes. The height medians are specified above the bars. The black dots indicate a posterior probability equal or above 0.95 and the grey dots show a posterior probability comprises between 0.90 and 0.95. The time scale is in million-year unit.

**Supplementary Figure S8:** **PC 3 and 4 of the Principal Component Analysis (PCA) at the species level.** (A) Individual factor map. (B) Variable factor map**.** The definitions of the CHELSA variables are given in the **supplementary Table S6**.

**Supplementary Figure S9: Principal Component Analysis (PCA)** **at the sub-species level.** (A) Biplot for *B. lunaria* subspecies showing PC 1 and 2. (B) Biplot for *B. lunaria* subspecies showing PC 2 and 3. (C) Biplot for *species 6* subspecies showing PC 1 and 2. (D) Biplot for *species 6* subspecies showing PC 2 and 3. (E) Biplot for *B. neolunaria* subspecies showing PC 1 and 2. (F) Biplot for *B. neolunaria* group subspecies showing PC 2 and 3. The definitions of the CHELSA variables are given in the **supplementary Table S6**.

*Supplementary Tables*

[See separate file, available from Zenodo: https://doi.org/10.5281/zenodo.6531504]

**Supplementary Table S1**: Information about the specimens used in this study.

**Supplementary Table S2**PCR protocols and thermocycling conditions developed in this study.

**Supplementary Table S3:** Information about the sequences used in the divergence time analyses at the Botrychioideae subfamily level.

**Supplementary Table S4**: Flow cytometry results.

**Supplementary Table S5**: Spore length measurements.

**Supplementary Table S6**: CHELSA climatic variable meanings.

*Supplementary Files*

[Available from Zenodo: https://doi.org/10.5281/zenodo.6531504]

**File S1**: Multiple alignment of the full dataset (A2) used for the phylogenetic reconstruction.

File name: File_S1_A2_full_alignment.zip

**File S2**: Multiple alignment of the reduced dataset (A1) used for the phylogenetic reconstruction.

File name: File_S2_A1_reduced_alignment.zip

**File S3**: Multiple alignment of the subfamily Botrychioideae used for the time divergence analyses.

File name: File_S3_time_calibration_Botrychioideae_alignment.zip

**File S4**: Multiple alignment A1 used for the time divergence analyses. This alignment is the alignment containing no duplicated sequenced produced by RAxML-NG.

File name: File_S4_time_calibration_B.lunaria_group_alignment.zip

**File S5**: Maximum likelihood tree inferred from A2 alignment with bootstrap support values.

File name: File_S5_ML_A2_12102020.zip

**File S6**: Bayesian tree inferred from A1 alignment with posterior probabilities.

File name: File_S6_BP_A1_15102020.zip

**File S7**: Maximum likelihood tree inferred from A1 alignment with bootstrap support values.

File name: File_S7_ML_A1_15102020.zip

**File S8**: Bayesian tree inferred from A1 alignment with indels scored. Contains the posterior probabilities.

File name: File_S8_BP_A1_indels_15102020.zip

**File S9**: Maximum likelihood tree inferred from A1 alignment with indels scored. Contains the bootstrap support values.

File name: File_S9_ML_A1_indels_15102020.zip

**File S10**: RAxML-NG bestTree of the phylogenetic tree inferred from A1 alignment (File S7) used to constraint the backbone of Maximum likelihood tree inferred from A2 alignment (File S5)

File name: File_S10_ML_A1_bestTree_15102020.zip

**Files S11-S23**: Summary of the Analysis of Variance, Tukey post-hoc tests results, Principal component analysis eigenvalues and variable contributions. Contained in Folder S1.

Folder name: Folder_S1_climatic_analyses.zip
